# The interplay between splicing of exons 18a and 47 differentially affects membrane targeting and function of human Ca_V_2.2

**DOI:** 10.1101/2023.09.15.557884

**Authors:** Shehrazade Dahimene, Karen M Page, Manuela Nieto-Rostro, Wendy S Pratt, Annette C Dolphin

**Author notes:** joint first authors (Shehrazade Dahimene, Karen M Page). Corresponding author: Annette C. Dolphin, Department of Neuroscience, Physiology and Pharmacology, University College London, Gower Street, London, WC1E 6BT, UK.

## Abstract

N-type calcium channels (Ca_V_2.2) are predominantly localized in presynaptic terminals, and are particularly important for pain transmission in the spinal cord. Furthermore, they have multiple isoforms, conferred by alternatively-spliced or cassette exons, which are differentially expressed. Here we have examined alternatively-spliced exon47 variants that encode a long or short C-terminus in human Ca_V_2.2. In the *Ensembl* database, all short exon47-containing transcripts were associated with the absence of exon 18a, therefore we also examined effect of inclusion or absence of exon18a, combinatorially with the exon47 splice variants. We found that long exon47, only in the additional presence of exon18a, results in Ca_V_2.2 currents that have a 3.6-fold greater maximum conductance than the other three combinations. In contrast, cell surface expression of Ca_V_2.2 in both tsA-201 cells and hippocampal neurons is increased ∼4-fold by long exon47 relative to short exon47, in either the presence or absence of exon18a. This surprising discrepancy between trafficking and function indicates that cell surface expression is enhanced by long exon47, independently of exon 18a. However, in the presence of exon47, exon18a mediates an additional permissive effect on Ca_V_2.2 gating. We also investigated the SNP in exon47 that has been linked to schizophrenia and Parkinson’s disease, which we found is only non-synonymous in the short exon47 C-terminal isoform, resulting in two minor alleles. This study highlights the importance of investigating the combinatorial effects of exon inclusion, rather than each in isolation, in order to increase our understanding of calcium channel function.

**Graphical abstract:** 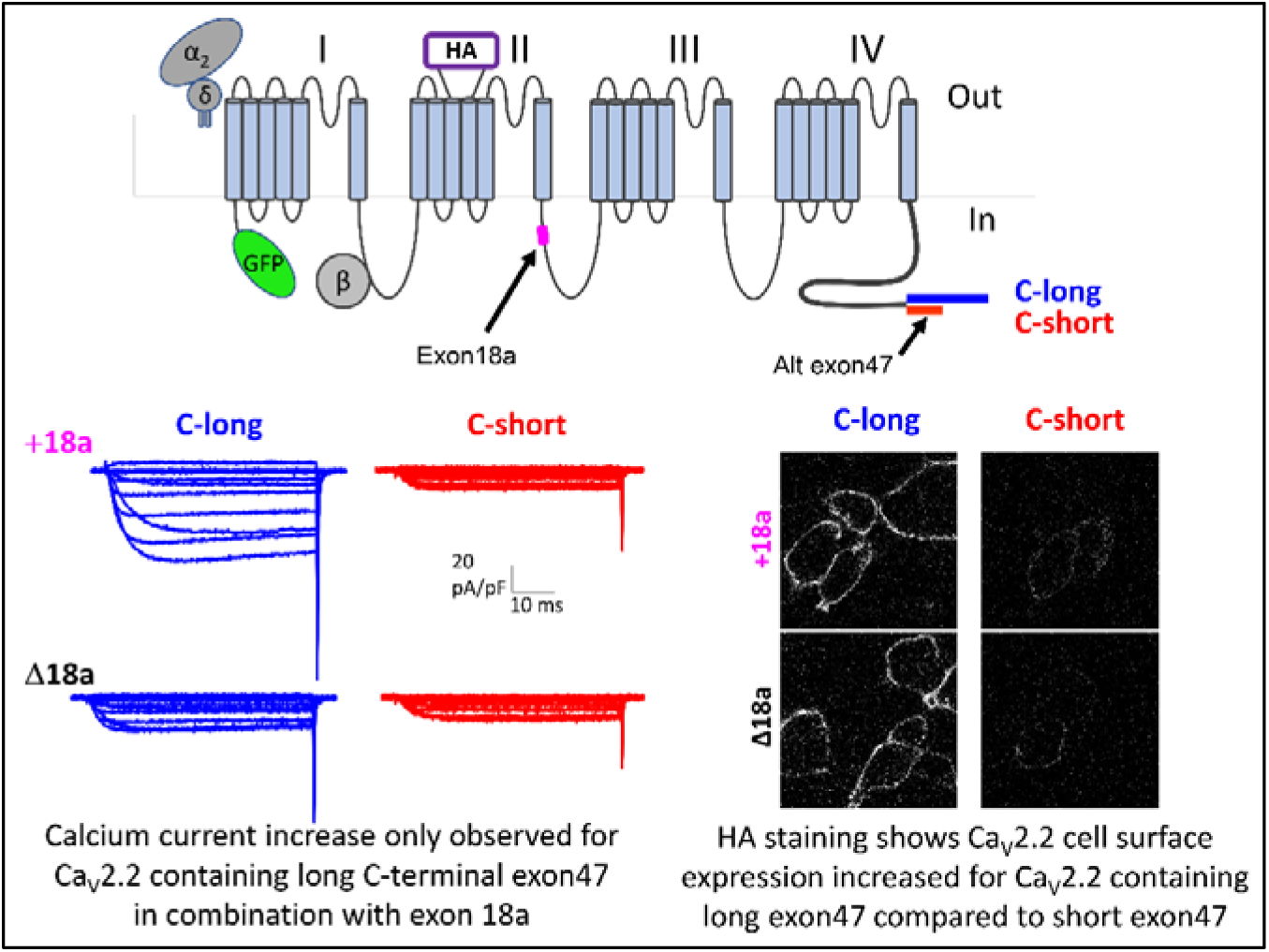

## Introduction

Synaptic transmission in the central and peripheral nervous system is dependent on voltage-gated calcium (Ca) channels that provide Ca^2+^ for vesicular release from presynaptic terminals. The particular channels involved are predominantly Ca_V_2.1 and Ca_V_2.2 ^1,2^. These channels are associated with auxiliary α_2_δ and β subunits which optimise their trafficking and function ^3-6^.

*CACNA1A*, the gene encoding Ca 2.1, is associated with multiple genetic disorders ^7,8^; in contrast there are very few identified human or animal pathological variants of *CACNA1B,* encoding Ca_V_2.2 which underlies N-type CaV channels ^9-11^. Nevertheless, Ca_V_2.2 knockout mice are associated with altered pain sensation and other phenotypes ^12^. These channels are particularly important for neurotransmission at primary afferents, including nociceptor terminals ^13,14^, and also in the sympathetic nervous system ^15^. A human variant in *CACNA1B* was linked to myoclonus-dystonia syndrome, although this was then disputed ^10,11^. A recent study identified novel risk loci in a genome-wide association (GWAS) analysis of Parkinson’s disease and schizophrenia, including a single nucleotide polymorphism (SNP), Rs2278973, in *CACNA1B* ^16^. However, upon analysis, we identified that this SNP was only non-synonymous in an alternatively-spliced exon47, which encodes a shorter C-terminal sequence than long exon47 (Figure 1A, B). The predominant residue at this SNP is arginine (Arg), and the two minor SNPs encode leucine (Leu) and histidine (His) (Supplementary Figure 1).

**Figure 1.**
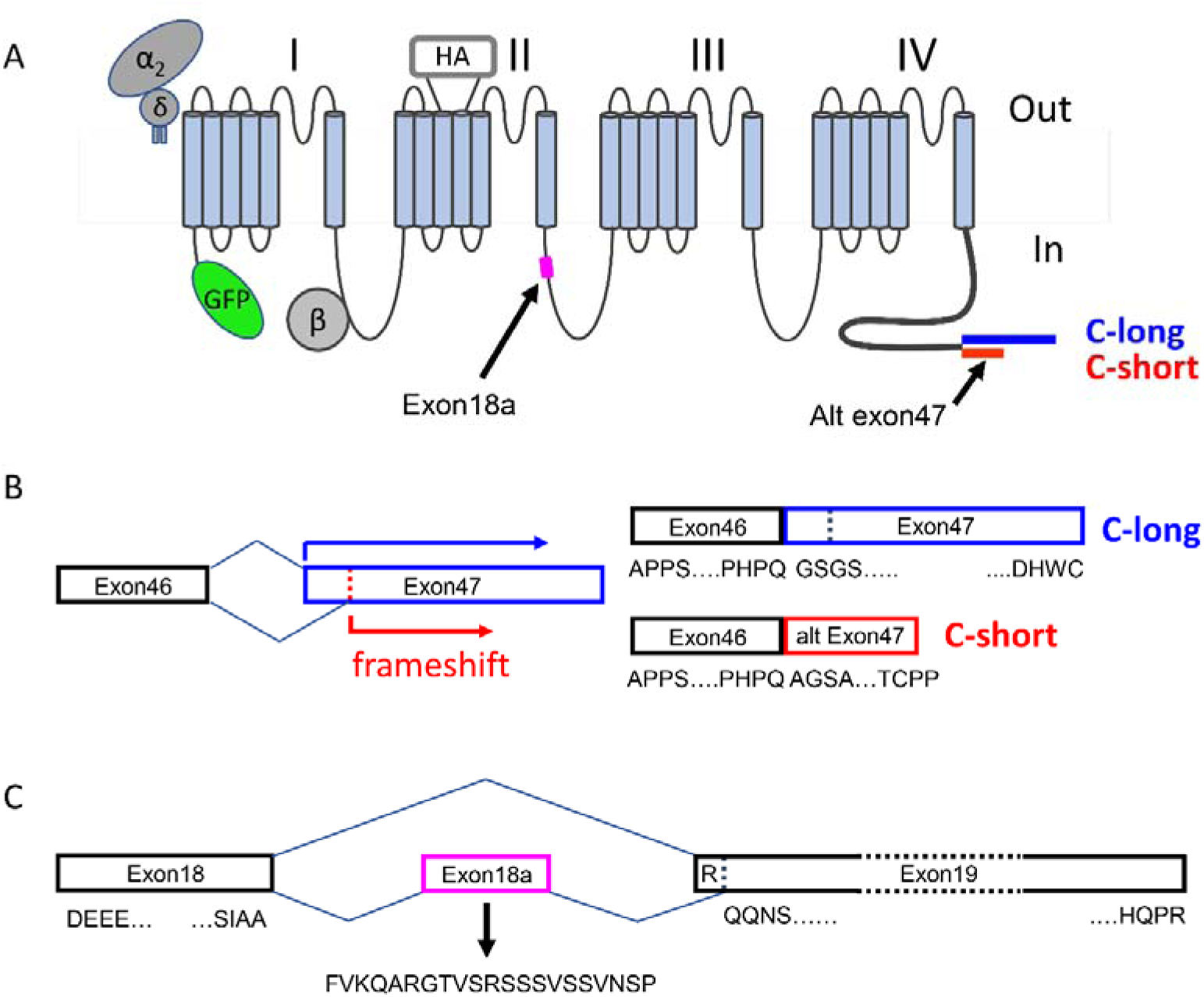
Organization of alternate exon47 and cassette exon18a in human Ca_V_2.2. **A** Schematic diagram of Ca 2.2 showing the GFP tag on the N terminus and the HA tag in the second extracellular loop of domain II. The position of alternatively-spliced exon18a is shown in the II-III loop and alternatively-spliced exon47, giving rise to the variant with the long C terminus or short C terminus is shown as the final exon at the end of the C terminus. **B** An alternative splice site within exon47 leads to a frameshift, resulting in an alternatively-spliced variant (labelled C-short, red). Long exon47 is shown in blue (C-long). Amino acid sequences at the beginning and end of the exons are indicated below. The full DNA sequence, together with translations for the long and short variants, is shown in Supplementary Figure 1. **C** Constitutively spliced exons 18 and 19 are shown in black. Amino acid sequences at the start and end of these exons are shown below. Alternatively-spliced exon18a is shown in pink, with the additional 21 amino acids indicated below. In the absence of exon18a, exon 19 begins with an R, which is missing in the presence of exon18a.

We further noted from a BLAST search (NCBI, *Ensembl ^17^*) of the short exon47 splice variant that all reported human Ca_V_2.2 sequences containing this short exon47 variant were also associated with absence of cassette exon18a (Figure 1C), whereas exon18a appeared to undergo alternative splicing in the Ca_V_2.2 variant with the long C terminus.

It has previously been reported that the inclusion of exon18a, which encodes a 21 amino acid sequence in the intracellular linker connecting Domains II and III of Ca_V_2.2, resulted in two-fold increase in Ca_V_2.2/β3 currents, when these channel combinations were expressed in a cell line ^18^. The mechanism controlling insertion of exon18a was found to involve the RNA binding protein Rbfox2, which suppressed exon18a inclusion ^18^. In further support of the effect of exon18a inclusion, ω-conotoxin GVIA-sensitive N-type Ca_V_ currents were larger in sympathetic neurons from mice engineered to express exon18a, compared to those not expressing it ^18^.

In the present study, we first examined the effect of inclusion of the long or short exon47 variants on Ca_V_2.2 expression, function and trafficking. Since our analysis suggests there is differential expression of exon18a associated with the exon47-containing isoforms, we also examined effect of the inclusion or absence of exon18a combinatorially with the two exon47 splice variants. We then determined whether there was any effect of the SNPs in short exon47 of*CACNA1B* ^16^.

Our results reveal a surprising discrepancy between Ca_V_2.2 trafficking and function. We show that Ca_V_2.2 containing the long exon47, in conjunction with exon18a, supports a large increase in Ca_V_2.2 current amplitude, whereas long exon47 mediates an increase in Ca_V_2.2 trafficking in tsA-201 cells and hippocampal neurons, irrespective of the presence or absence of exon18a.

## Methods

### Cell culture

#### tsA-201 cells

The tsA-201 cells (European Collection of Cell Cultures, female sex) were plated onto cell culture flasks or coverslips coated in poly-L-lysine, and cultured in Dulbecco’s Modified Eagle Medium (DMEM) supplemented with fetal bovine serum (5 %), Penicillin-Streptomycin (1 %) and GlutaMAX (Thermo Fisher Scientific, 1 %), in a 5 % CO2 incubator at 37 °C.

#### Hippocampal neurons

Tissue was obtained from P0/P1 rat pups (Sprague-Dawley, both sexes). All experiments were performed in accordance with the UK Home Office Animals (Scientific procedures) Act 1986, with UCL ethical approval. Briefly, hippocampi were dissected, chopped into small pieces and treated for 40 min at 37 °C with a papain solution containing: 7 units /ml of papain, 0.2 mg/ml L-cysteine, 0.2 mg/ml bovine serum albumin (BSA), 1000 units/ml DNase-1 and 5 mg/ml glucose (all from Sigma Aldrich) in Hank’s basal salt solution medium (Thermo Fisher Scientific). Hippocampi were then washed with inactivation medium (Minimum Essential Medium (MEM); Thermo Fisher Scientific, 5% FBS, 0.38 % glucose, 0.25% BSA) and mechanically dissociated in serum medium (MEM, 5 % FBS, 1.38 % glucose). After centrifugation at 1000 rpm for 10 min, cells were resuspended in serum medium and seeded onto coverslips precoated with poly-D-lysine hydrobromide (Sigma; 50 µg/ml). 1 h later, cells were covered with serum-free neuronal plating medium comprising Neurobasal Medium, supplemented with B27 (2 %), GlutaMAX (1 %) (all from Thermo Fisher Scientific). Neurons were kept in a 5 % CO2 incubator at 37 °C and half the medium was replaced every 3-4 days. At 7 days in vitro and 2 h before transfection, half of the medium was removed, and fresh medium was added.

### Expression constructs and mutagenesis

The following expression constructs were used: human Ca_V_2.2 (huCaV2.2 (pSAD442-1) was a gift from Diane Lipscombe (Addgene plasmid # 62574; http://n2t.net/addgene:62574; RRID:Addgene_62574)), from which were derived GFP_Ca_V_2.2-HA-C-long, GFP_Ca_V_2.2-HA-C-short, GFP_Ca_V_2.2-HA-C-long(Δexon18a), GFP_Ca_V_2.2-HA-C-short(Δexon18a), GFP_Ca_V_2.2-HA-C-short-Arg and GFP_Ca_V_2.2-HA-C-short-His. Other constructs used were β1b (rat, X61394) ^19^, α_2_δ-1 (rat, M86621), and the transfection markers, mCherry ^20^ and CD8 ^21^ where stated. A double-HA tag was added to an extracellular loop in domain II of the human Ca_V_2.2 in the same position as used previously in the rabbit Ca_V_2.2 (Cassidy, Ferron et al. 2014) and GFP was added to the N-terminus. All the constructs were verified by Sanger sequencing (Source Bioscience) and have been submitted to Addgene. All cDNAs were used in either pcDNA3 vector for expression in tsA-201 cells or pCAGGS for hippocampal neurons.

### Antibodies

The following antibodies were used: anti-hemagglutinin (HA, rat, Sigma-Aldrich). For immunoblotting, secondary goat anti-rat Horseradish Peroxidase (HRP; Biorad) was used. For immunocytochemistry, secondary anti-rat-Alexa Fluor 594 and anti-rat-Alexa Fluor 647 antibodies (1:500, Life Technologies) were used to visualise the HA-tagged channels.

### Cell transfections

#### tsA-201 cells

The tsA-201 cells were transfected with PolyJet in a 3:1 ratio with DNA mix, according to the manufacturer’s protocol. Following incubation overnight at 37 °C, culture medium was changed. The transfection mix consisted of plasmids containing cDNAs encoding each of the GFP_Ca_V_2.2-HA variants, β1b and α_2_δ-1 in a ratio of 3:2:2. For electrophysiological experiments, CD8 was used as a transfection marker (ratio 3:2:2:1).

#### Hippocampal neurons

The transfection mix consisted of plasmids containing cDNAs encoding the GFP_Ca_V_2.2-HA variants, β1b, α_2_δ-1 and mCherry in a ratio of 3:2:2:0.3. Neurons were transfected with Lipofectamine 2000 (Life Technology), according to the manufacturer’s protocol with 1 μl Lipofectamine and 2 μl DNA mix per sample.

### Electrophysiology

Ca_V_2.2 currents in transfected tsA-201 cells were investigated by whole cell patch clamp recording. The patch pipette solution contained in mM: Cs-aspartate, 140; EGTA, 5; MgCl_2_, 2; CaCl_2_, 0.1; K_2_ATP, 2; HEPES, 10; pH 7.2, 310 mOsm with sucrose. The external solution for recording Ba^2+^ currents contained in mM: tetraethylammonium (TEA) Br, 160; KCl, 3; NaHCO_3_, 1.0; MgCl_2_, 1.0; HEPES, 10; glucose, 4; BaCl_2_, 1, pH 7.4, 320 mosM with sucrose. 1 mM extracellular Ba^2+^ was the charge carrier. Pipettes of resistance 2–4 MΩ were used. An Axopatch 1D or Axon 200B amplifier was used, and whole cell voltage-clamp recordings were sampled at 10 kHz frequency, filtered at 2 kHz and digitized at 1 kHz. 70–80% series resistance compensation was applied and all recorded currents were leak-subtracted using P/8 protocol. Analysis was performed using Pclamp 9 (Molecular Devices) and Origin 7 (Microcal Origin, Northampton, MA). To obtain the maximum conductance (*G_max_*), Current-voltage (*IV)* relationships were fitted by a modified Boltzmann equation as follows: *I=G_max_*(V−V_rev_)/(1+exp(−(V−V_50, act_)/k))* where *I* is the current density (in pA/pF), *G_max_* is in nS/pF, *V_rev_* is the apparent reversal potential, *V_50, act_* is the midpoint voltage for current activation, and *k* is the slope factor.

### Immunoblotting

Immunoblotting was carried out on tsA-201 cells expressing the cDNAs as described. At 48 h after transfection, cells were rinsed with phosphate buffered saline (PBS, pH 7.4), scraped into cold PBS and centrifuged at 1,000 x *g* at 4⁰C for 10 minutes. Cell pellets were homogenised in PBS containing 1% Igepal, 0.1% SDS and protease inhibitors (PI, cOmplete, Sigma-Aldrich), pH 7.4, and then incubated on ice for 30 min to allow cell lysis. Whole cell lysates (WCL) were centrifuged at 20,000 x g for 25 min at 4⁰C and supernatants were assayed for total protein (Bradford assay, BioRad). Aliquots of WCL (30 µg total protein, per sample) were diluted with 2x Laemmli buffer supplemented with 100 mM dithiothreitol and incubated at 60°C for 10 min ^22^. Samples were resolved by SDS-PAGE on 3%–8% Tris-Acetate gels (Thermo Fisher Scientific) and transferred to polyvinylidene fluoride (PVDF) membranes (Biorad). Membranes were blocked with 3% bovine serum albumin (BSA), 0.5% Igepal in Tris-buffered saline (TBS, pH 7.4) for 30 min at room temperature and then incubated overnight at 4°C with anti-HA (rat, 1:1000 dilution, Sigma-Aldrich) antibody in TBS containing 10% goat serum, 3% BSA and 0.02% Igepal (pH 7.4). After washing in TBS containing 0.5% Igepal, membranes were incubated with the secondary antibody, goat anti-rat Horseradish Peroxidase (HRP, 1:2000 dilution, Biorad) for 1 h at room temperature. The signal was obtained using the ECL 2 assay (Thermo Scientific) and membranes were scanned on a Typhoon 9410 phosphorimager (GE Healthcare).

### Immunocytochemistry

Cells were fixed with 4 % paraformaldehyde in PBS for 5 min, incubated with blocking buffer (20 % goat serum, 4 % BSA in PBS) for 1 h at room temperature before being incubated with rat anti-HA (Roche) diluted either 1:250 (tsA-201cells) or 1:100 (hippocampal neurons) in 0.5 x blocking buffer for 1 h at room temperature (tsA-201 cells) or at 4 °C overnight (hippocampal neurons). After washing, samples were incubated with secondary antibodies, anti-rat Alexa Fluor 594 (tsA cells) or anti-rat Alexa Fluor 647 (hippocampal neurons) at a dilution of 1:500 for 1 h at room temperature. Coverslips were washed, stained with 0.5 µM 4’,6’-diamidino-2-phenylindole (DAPI), and mounted in VectaShield (Vector Laboratories). Imaging was performed on Zeiss LSM 780 confocal microscope.

### Image analysis

The tsA-201 cell images were obtained using a 63 × objective at a resolution of 1024 × 1024 pixels and an optical section of 0.5 μm. After choosing a region of interest containing transfected cells, the 3 × 3 tile function of the microscope was used to allow a larger area to be selected without bias. Images were then analyzed using Fiji. Every transfected cell was included. Surface labelling (HA staining) was measured using a freehand line tool of 5 pixels (0.65 μm) width and manually tracing the surface of the cells. Intracellular GFP staining was measured by drawing around the cell (omitting the nucleus and the plasma membrane). The value of the mean pixel intensity in different channels was measured separately and background was subtracted by measuring the intensity of an imaged area without transfected cells.

Hippocampal neurons were imaged using a 20 × objective with a 5 μm optical section. The fluorescence intensity along neuronal projections was assessed as follows: two concentric circles of 100 μm and 150 μm diameter were drawn around each neuronal cell body. A freehand line tool of 3 μm width tracing the neuronal processes (3 to 5 per neuron) between the circles was drawn in the mCherry images and used as template for GFP and HA images. Hippocampal somata were imaged at 63 x objective with a 1 μm optical section. For each neuron selected as positive for mCherry, immunofluorescence (HA and GFP) was measured by two ROIs, one ROI manually traced around the cell body, using a freehand line tool (0.4 μm width) to analyse the cell surface staining and another ROI drawn between the cell surface and the nucleus to determine the intracellular staining. Total staining was calculated per each cell combining the 2 ROIs. The mean pixel intensity in the different channels were measured and the background was subtracted from each image.

### Data analysis and statistics

Quantitative data are presented by GraphPad Prism 8 software (v8.0.0; GraphPad Software, Inc.) or Origin-Pro 2021 as the mean ± standard error of the mean (SEM) and individual data points as described. One-way analysis of variance (ANOVA) followed by a Šidák *post hoc* test for multiple comparisons were used.

## Results

In this study we have used human Ca_V_2.2 containing the following alternatively-spliced exons: +exon 10a, +exon18a, Δexon24a, +exon31a, +exon37b and +long exon47 ^10^. We added an extracellular HA tag (two HA sequences separated by a Gly residue) in an extracellular linker of domain II, in the corresponding position to that in the rabbit Ca_V_2.2 we have used previously ^6^, and a GFP tag to the N terminus ^23,24^ (Figure 1A), to form GFP_Ca_V_2.2-HA-C-long. It contains the alternatively-spliced 21 amino acid exon18a in the II-III linker (Figure 1C). We used this construct to generate GFP_Ca_V_2.2-HA-C-short, which uses an alternative splice site part-way through exon47 (Supplementary Figure 1). This creates a frame-shift, resulting in an alternative, shorter C terminus (Figure 1B). The sequence for exon18a was then deleted from both these constructs to give GFP_Ca_V_2.2-HA-C-long-Δexon18a and GFP_Ca_V_2.2-HA-C-short-Δexon18a.

### The combination of C-terminal long exon47 together with exon18a in the II-III linker results in an increase in Ca_V_2.2 currents

We next examined the currents generated by the GFP_Ca_V_2.2-HA constructs expressed in tsA-201 cells, together with β1b and α_2_δ-1 subunits. For GFP_Ca_V_2.2-HA-C-long, the additional presence of exon18a resulted in a ∼3.6-fold increase of the maximum conductance (G_max_), compared to the absence of exon18a (Figure 2A-C). Surprisingly, for GFP_Ca_V_2.2-HA-C-short (containing short exon47), the inclusion of exon18a produced no increase in calcium currents (Figure 2A-C), which were of very similar magnitude to those produced by GFP_Ca_V_2.2-HA-C-long-Δexon18a.

**Figure 2.**
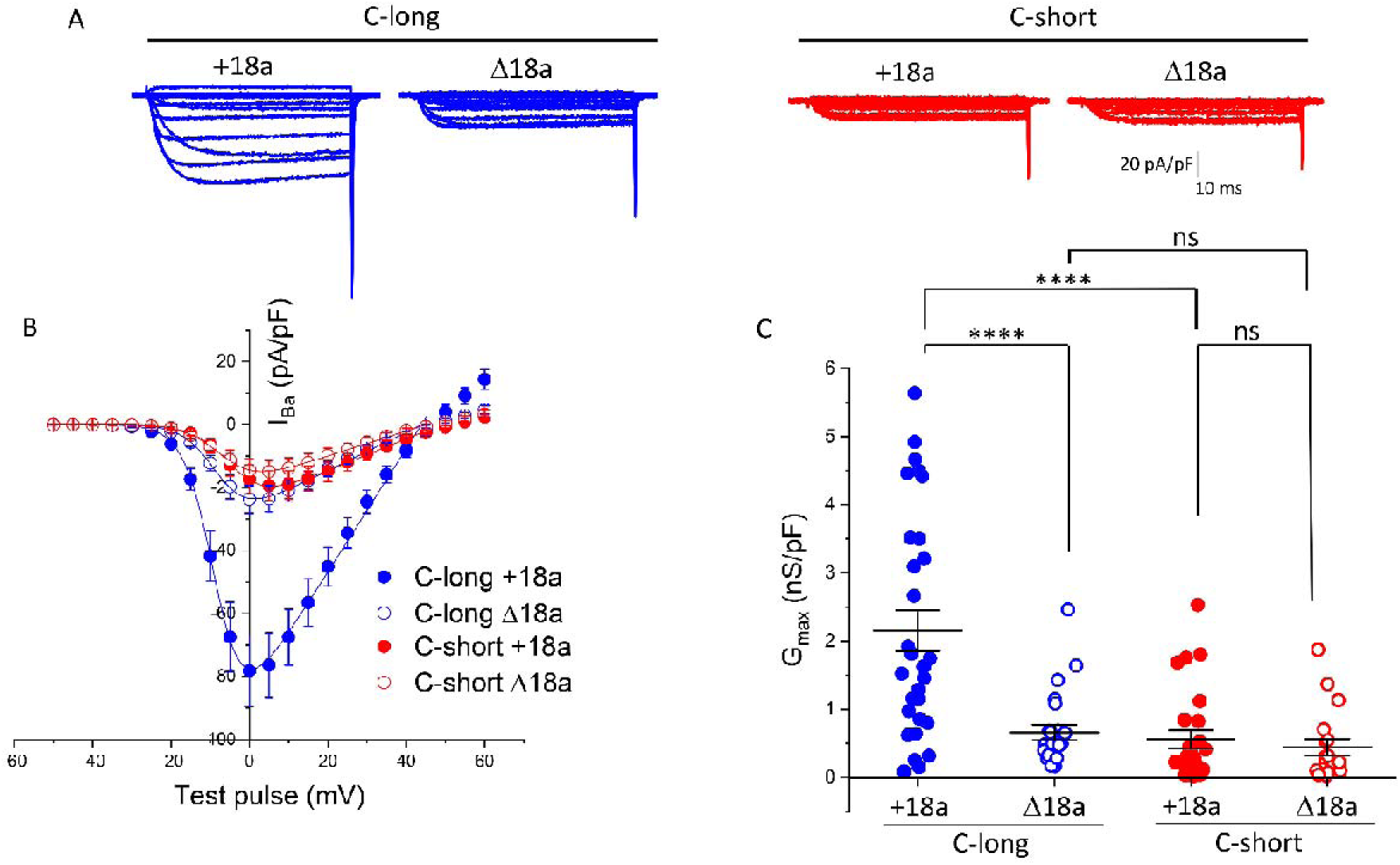
Long exon47 increases Ca_V_2.2 currents, only in the presence of exon18a. **A** Example of whole-cell patch-clamp recordings for human GFP_Cav2.2-HA having the long (C-long) or short (C-short) variant of exon47 in the presence (+18a) or absence (Δ18a) of alternatively-spliced exon18a. All conditions are in the presence of α_2_δ-1 and β1b. Holding potential −80 mV, steps between −50 and +60 mV for 50 ms (applies to all traces). **B** Mean (± SEM) *IV* relationships for the conditions shown in A. GFP_Ca_V_2.2-HA, C-long +18a (*n*⍰=⍰30, blue solid circles), C-long Δ18a (*n*⍰=⍰23, blue open circles), C-short +18a (*n*⍰=⍰18, red solid circles) and C-short Δ18a (*n*⍰=⍰26, red open circles). The individual and mean data were fit with a modified Boltzmann equation (see Materials and Methods). The potential for half-maximal activation (V_50,act_) (mV) was -6.07 ± 0.76, -7.51 ± 0.71, -4.1± 0.73 and -5.13 ± 1.04 for C-long +18a, C-long Δ18a, C-short +18a and C-short Δ18a respectively. **C** G_max_ (nS/pF)from the *IV* relationships shown in B. Individual data (same symbols as in B) and mean⍰±⍰SEM are plotted. Statistical significance was determined using one-way ANOVA followed by Šidák’s *post hoc* test for multiple comparisons. ****P⍰<⍰0.0001, ns: non-significant.

### The presence of C-terminal long exon47 is important for cell surface expression of Ca_V_2.2

In order to determine whether the marked increase in Ca_V_2.2 current amplitude observed for the long exon47 variant of Ca_V_2.2, in conjunction with the presence of exon18a, was due to an increase in cell surface expression for this exon combination, we next expressed the GFP_Ca_V_2.2-HA constructs, together with β1b and α_2_δ-1 subunits, in tsA-201 cells and used confocal microscopy to visualize expression of the GFP_Ca_V_2.2-HA subunit at the cell surface. Transfected cells were identified by expression of the GFP tag on GFP_Ca_V_2.2-HA. Ca_V_2.2 on the plasma membrane was measured by anti-HA antibody binding to the extracellular HA tag of GFP_Ca_V_2.2-HA, in non-permeabilised conditions. We found that cell-surface expression of the GFP_Ca_V_2.2-HA-C-long(+exon18a) variant was increased by 4.5-fold compared with GFP_Ca_V_2.2-HA-C-short(+exon18a) (Figure 3A, B i). However, in contrast to the calcium current measurements (Figure 2), deletion of exon18a had no significant effect on HA expression at the plasma membrane for either the C-long or C-short Ca_V_2.2 variants. Cell-surface expression of the GFP_Ca_V_2.2-HA-C-long(Δexon18a) variant was still increased by 3.2-fold compared with GFP_Ca_V_2.2-HA-C-short(Δexon18a) (Figure 3A, B i).

**Figure 3.**
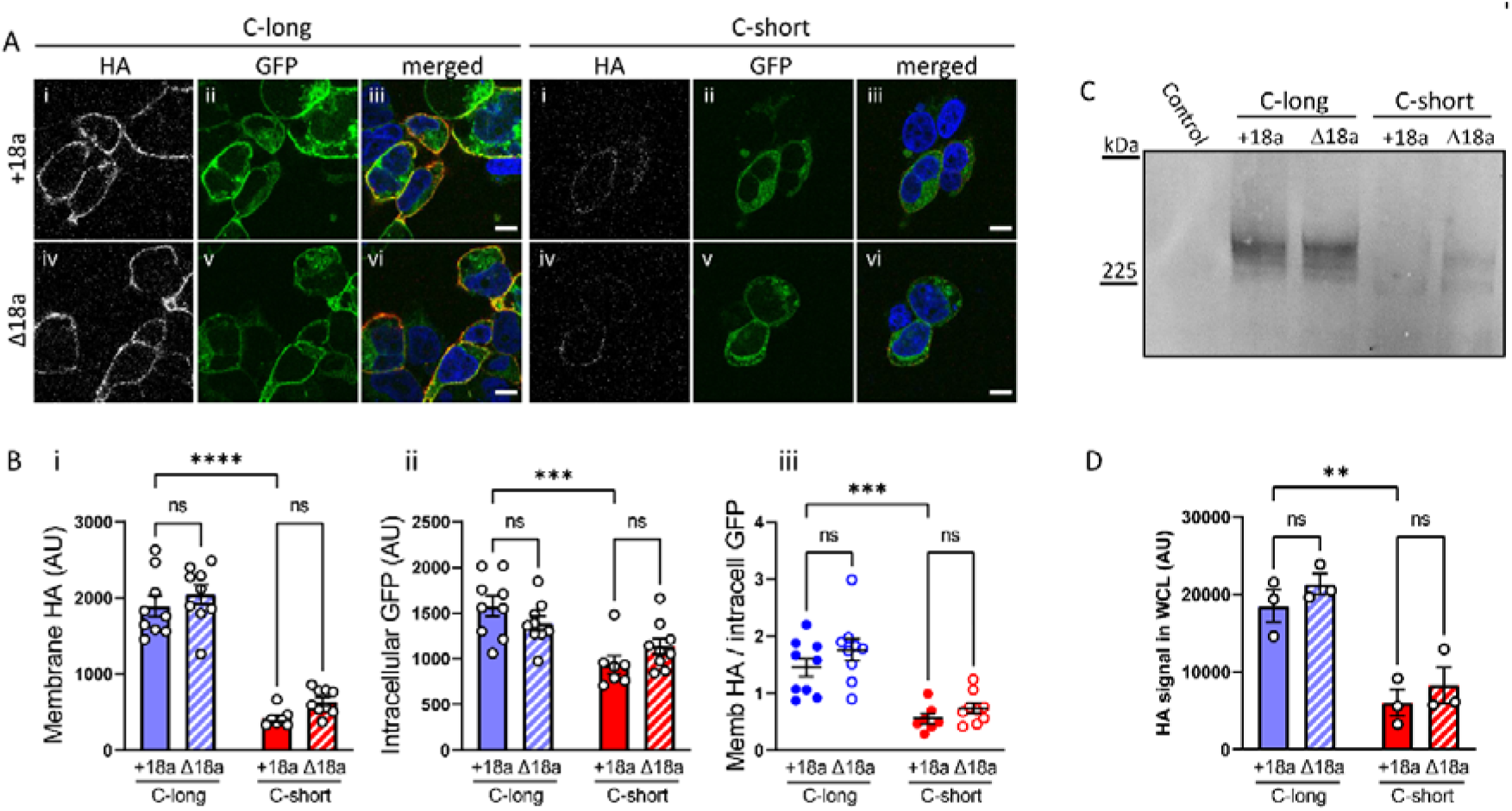
Long exon47 increases cell surface expression of Ca_V_2.2 irrespective of exon18a inclusion. **A** Representative tsA-201 cells expressing GFP-Ca_V_2.2-HA having the long (left) or short (right) variant of exon47 in the presence (top row, +18a) or absence (bottom row, Δ18a) of alternatively-spliced exon18a. All conditions are in the presence of α_2_δ-1 and β1b. Cells were incubated with anti-HA antibody in non-permeablised conditions to show HA staining on the extracellular side of the plasma membrane (panels i and iv), to be compared with intracellular GFP fluorescence (ii and v). Merged images (nuclei stained with DAPI in blue and HA in red) are shown in panels iii and vi. Scale bar = 10 μm. **B** Quantification of HA staining at the plasma membrane (i) or intracellular GFP fluorescence (excluding the nucleus and plasma membrane, ii), showing mean ± SEM for GFP-CaV2.2-HA C-long containing exon18a (blue solid) or without exon18a (blue striped), C-short with exon18a (red solid) or without exon18a (red striped). Individual data points show the mean of 30-100 cells from 7-9 different transfections in 3 independent experiments. The ratio of HA/intracellular GFP for each individual cell was calculated; mean ± SEM for different transfections is shown in iii. Statistical significance was determined using one-way ANOVA followed by a Šidák *post hoc* test for multiple comparisons: **** p<0.0001, *** p<0.001, ns: non-significant. **C** Immunoblot of whole-cell lysates (WCL) from tsA-201 cells transfected with either GFP_Ca_V_2.2-HA C-long or C-short in the presence (+18a) or absence (Δ18a) of alternatively-spliced exon18a together with α_2_δ-1 and β1b. The immunoblot was performed using an anti-HA antibody. Untransfected tsA-201 cells were used as control. **D** Mean⍰±⍰SEM and individual data-points of three separate experiments, including that in (C) showing quantification of the HA immunoblots. Statistical significance was determined using one-way ANOVA followed by a Šidák *post hoc* test for multiple comparisons. ** p<0.01, ns: non-significant.

There was a smaller increase in intracellular GFP_Ca_V_2.2-HA, as determined by GFP signal associated with the presence of C-long exon47, compared to C-short exon47 (1.7-fold in the presence of exon18a and 1.2-fold in its absence) (Figure 3A, B ii). From the ratio of cell surface/intracellular expression of the four Ca_V_2.2 variants, there was only a significant effect of the long exon47 to increase trafficking to the cell surface, and no effect of exon18a (Figure 3B iii).

These results suggest that the presence or absence of exon18a does not affect cell surface expression or trafficking of GFP_Ca_V_2.2-HA in tsA-201 cells but that trafficking to the plasma membrane is highly influenced by the presence of the long exon47 variant in the C-terminus.

We then performed immunoblotting with these constructs to determine whether there were similar effects on total Ca_V_2.2 protein expression in the tsA-201 cells (Figure 3C, Supplementary Figure 2). This showed that the level of full length GFP_Ca_V_2.2-HA-C-long protein was more than 2-fold greater than that of GFP_Ca_V_2.2-HA-C-short, regardless of the inclusion or absence of exon18a (Figure 3D), in a clear parallel with the immunocytochemistry results. Together these data indicate that the presence of short exon47 in Ca_V_2.2 confers a defect in cell surface trafficking relative to long exon47, and a reduction in the total full-length channel level.

### The presence of long exon47 is important for trafficking of Ca_V_2.2 in hippocampal neurons

We then examined the effect of both exon47 splice variants and the inclusion of exon18a on Ca_V_2.2 expression in hippocampal somata and neurites. In cell bodies, cell-surface expression of the GFP_Ca_V_2.2-HA-C-long (+exon18a) variant was increased by 4.3-fold compared with GFP_Ca_V_2.2-HA-C-short(+exon18a) (Figure 4A, B i). For the same pair of constructs there was a smaller increase (1.9-fold) in total Ca_V_2.2 expression attributable to long exon47, measured by GFP signal (Figure 4A, B ii). In contrast with the data in tsA-201 cells, the inclusion of exon18a also produced a small increase of HA expression at the plasma membrane (1.4-fold increase) and GFP signal (1.3-fold increase) for the long exon47 variant, but it had no effect for the short exon47 variant (Figure 4A, B i and B ii). However, from the ratio of cell surface/total expression of Ca_V_2.2, there was only a significant effect of the long exon47 to increase trafficking to the cell surface, and no effect of the inclusion of exon18a (Figure 4B iii).

**Figure 4.**
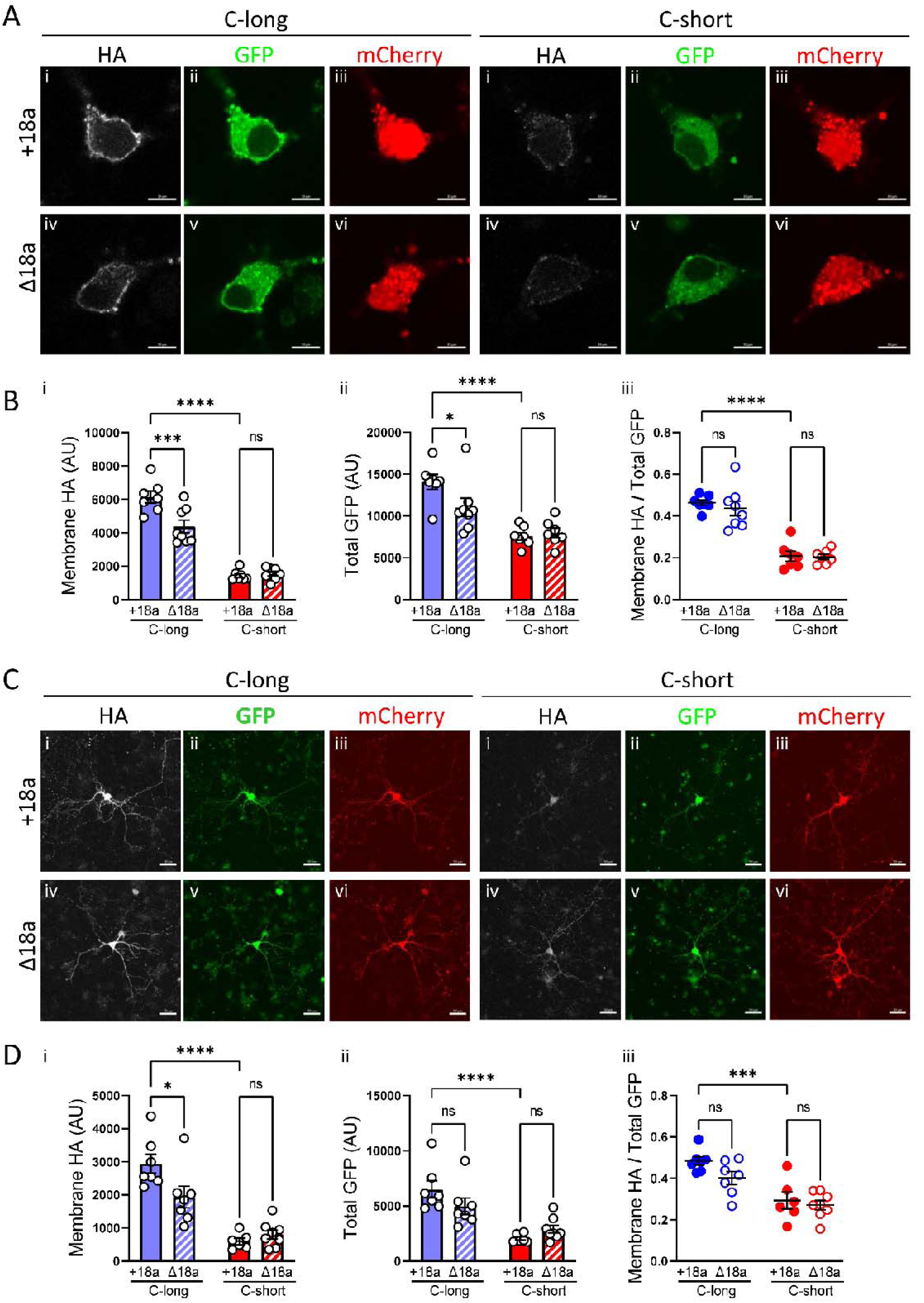
Effects of alternate C-terminal exon47 with or without exon18a on Ca_V_2.2 expression in soma and neurites of hippocampal neurons. **A, C** High (A) or low (C) magnification confocal images of hippocampal neurons expressing either the long or short C-terminal splice variants of GFP_Ca_V_2.2_HA with or without exon18a (+18a or Δ18a respectively), together with α_2_δ-1 and β1b. Scale bar 10 μm in A and 50 μm in C. Transfected neurons identified by the expression of mCherry in red (A, C iii and vi), also expressed GFP_Ca_V_2.2_HA that can be detected throughout the cell with GFP in green (A, C ii and v) and at the cell surface by immunolabelling with HA in white (i and iv). **B, D** Quantification of Ca_V_2.2 at the membrane (HA in B, D i) or total (GFP in B, D ii) and the ratio of HA/GFP (B, D iii) in the soma or projections of hippocampal neurons is shown in B and D respectively, where the bars show mean ± SEM, and each dot represents the mean value of a coverslip. In B, *n*=7 coverslips for all except for C-long Δ18a where *n*=8. The mean of each coverslips was calculated from 5 to 16 cells per coverslip. In D, *n*=7 coverslips for C-long +18a and Δ18a, *n*=6 for C-short +18a and *n*=8 for C-short Δ18a, where the mean of each coverslip was calculated from 6 to 10 neurons. Statistical significance is indicated with * P < 0.05, ** P<0.01, *** P<0.001, **** P<0.0001 and ns: non-significant (one-way ANOVA followed by Sidak’s multiple comparisons test).

Very similar results were observed in the neurites of these hippocampal neurons, in that the presence of the long exon47 variant induced a large increase (4.9-fold) of cell surface Ca_V_2.2 in the neurites relative to the short exon47 variant (Figure 4C, D i). There was also an increase (3.1-fold increase) in total Ca_V_2.2 expression for the long exon47 variant compared to the short exon47 variant, measured by GFP signal (Figure 4C, D ii). The presence of exon18a also induced a small increase in membrane expression of Ca_V_2.2 in the neurites, but only for the long exon47 variant (Figure 4D i). However, from the ratio of cell surface/total expression of Ca_V_2.2, there was only a significant effect of the long exon47, but not of exon 18a, to increase trafficking to the cell surface of the neurites (Figure 4D iii).

### Examination of the effect of SNP variants in the short exon47 of Ca_V_2.2

The Rs2278973 SNP gives no change in amino acid in the long exon47 variant of Ca_V_2.2. In the short exon47 variant, however, this SNP results in three alternative amino acids, Arg, Leu or His, at this position. Unlike the reference gene, *Ensembl* ENSG00000148408 (NCBI Reference Sequence: NM_000718.4), which would produce a Leu residue at this position, the Addgene plasmid #62574 that we used produces an Arg. Arg has a frequency of 91.7% in the population and, from comparison with other species, appears to be the ancestral gene (gnomAD genomes v3.1.2 database (*Ensembl*) https://www.ensembl.org/Homo_sapiens/Variation/Population?db=core;r=9:138121310-138122310;v=rs2278973;vdb=variation;vf=729552325).

We created two additional constructs in the GFP_Ca_V_2.2-HA-C-short (which contains exon18a and has an Arg at position 2236 of Ca_V_2.2, see Supplementary Figure 1), to give GFP_Ca_V_2.2-HA-C-short-Leu and GFP_Ca_V_2.2-HA-C-short-His, corresponding to the two minor alleles, which have population frequencies of 8.27% and 0.003% respectively.

Expression of these constructs, together with β1b and α_2_δ-1, in tsA-201 cells revealed that the HA signal at the plasma membrane was not significantly different for any of the amino acid substitutions in the short C-terminus (Figure 5A, B). Plasma membrane expression of GFP_Ca_V_2.2-HA-C-long was markedly increased when compared to all the C-short constructs (by 4-fold compared to GFP_Ca_V_2.2-HA-C-short-Arg, and by 4.6-fold compared to both the Leu and His-containing variants). Intracellular GFP expression of GFP_Ca_V_2.2-HA-C-long was increased by a smaller amount (1.7-fold) compared to all C-short variants (Figure 5C). This result was paralleled by an increase in the total GFP_Ca_V_2.2-HA protein level, determined by immunoblotting for GFP_Ca_V_2.2-HA-C-long, relative to all the C-short variants (Figure 5D, E).

**Figure 5.**
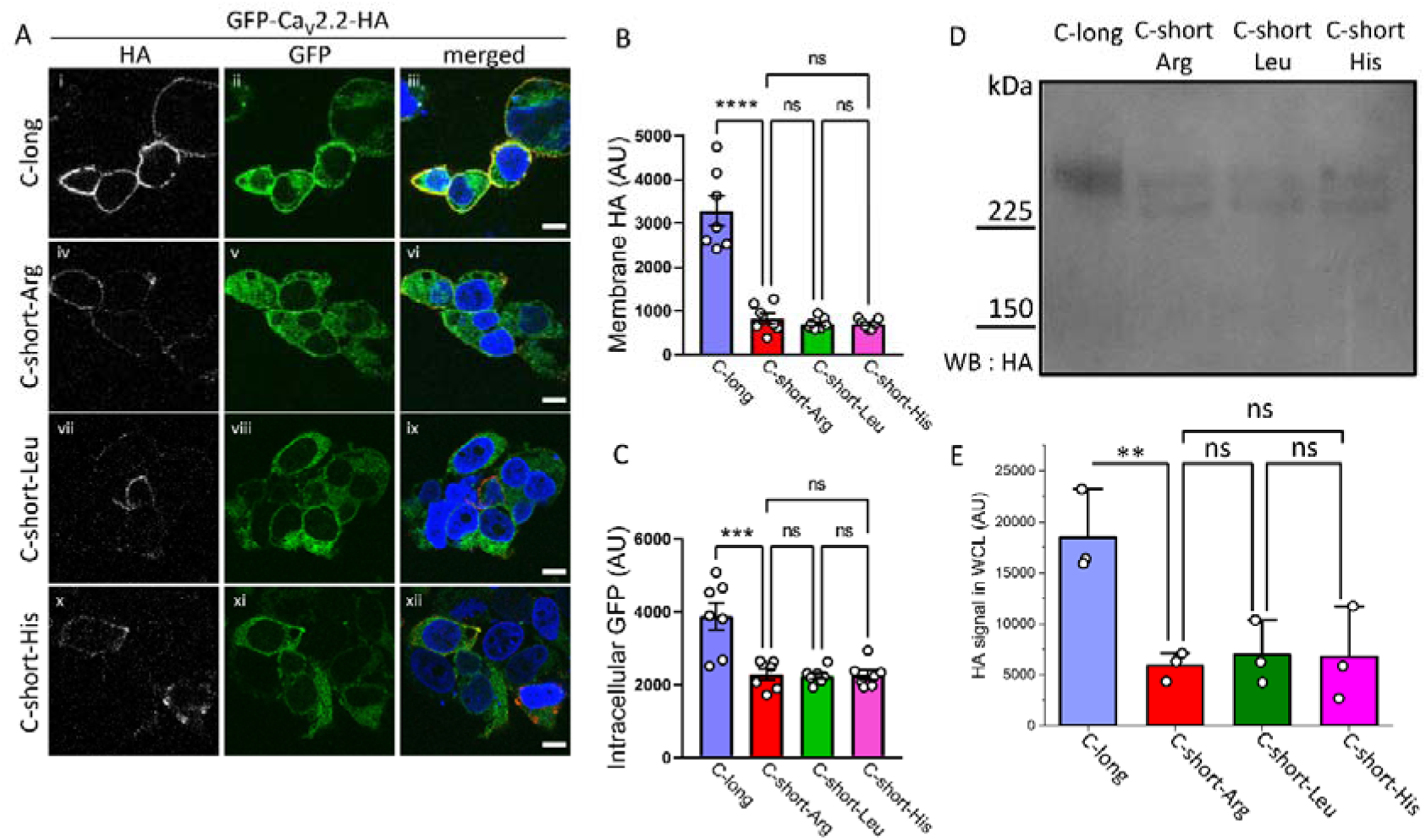
Non-synonymous SNPs in short exon47 do not influence cell surface or total expression of Ca_V_2.2. **A** Representative tsA-201 cells expressing GFP-Ca_V_2.2-HA having the long (i-iii) or short-Arg ancestral (iv-vi), short-Leu (vii-ix) or short-His (x-xii) Rs2278973 variants of C-short-exon47. All variants contain exon18a and are in the presence of α_2_δ-1 and β1b. Cells were incubated with anti-HA antibody in non-permeablised conditions to show HA staining on the extracellular side of the plasma membrane (white, left), to be compared with intracellular GFP fluorescence (green, centre). Merged images (nuclei stained with DAPI in blue and HA in red) are shown in the right panel. Scale bar = 10 μm. **B, C** Quantification of HA staining at the plasma membrane (B) or intracellular GFP fluorescence (excluding the nucleus and plasma membrane, C), showing mean ± SEM for GFP-Ca_V_2.2-HA C-long (blue), C-short (Arg, ancestral, red), C-short-Leu (green) or C-short-His (pink) variants. Individual data points show the mean of 30-100 cells from 7 different transfections in 2 independent experiments. Statistical significances: **** p<0.0001, *** p<0.001, ns: non-significant (one-way ANOVA followed by a Šidák *post hoc* test for multiple comparisons). **D** Immunoblot of whole-cell lysates (WCL) from tsA-201 cells transfected with either GFP_Ca_V_2.2-HA C-long, C-short-Arg, C-short-Leu or C-short-His, together with α_2_δ-1 and β1b. Immunoblot was performed using an anti-HA antibody. Full immunoblot given in Supplementary Fig. 2. **E** Mean⍰±⍰SEM and individual data-points of three separate experiments, including that in (**D**), showing quantification of the HA immunoblots for Ca_V_2.2-HA C-long (blue), C-short -Arg (red), C-short-Leu (green) or C-short-His (pink) variants. Statistical significance was determined using one-way ANOVA followed by a Šidák post hoc test for multiple comparisons: ** p<0.01, ns: non-significant.

These GFP_Ca_V_2.2-HA SNP variants (containing exon18a), together with β1b and α_2_δ-1, were then expressed in tsA-201 cells, and Ca_V_2.2 currents were recorded to examine whether there was any effect of the different SNP variants (Figure 6A-C). The data for GFP_Ca_V_2.2-HA-C-long, and GFP_Ca_V_2.2-HA-C-short are repeated from Figure 2 for comparison, as all experiments were performed contiguously. In agreement with the cell surface expression data, there was no difference between the Ca_V_2.2 G_max_ or other biophysical properties examined for the GFP_Ca_V_2.2-HA-C-short constructs containing Arg, Leu or His (Figure 6A-C).

**Figure 6.**
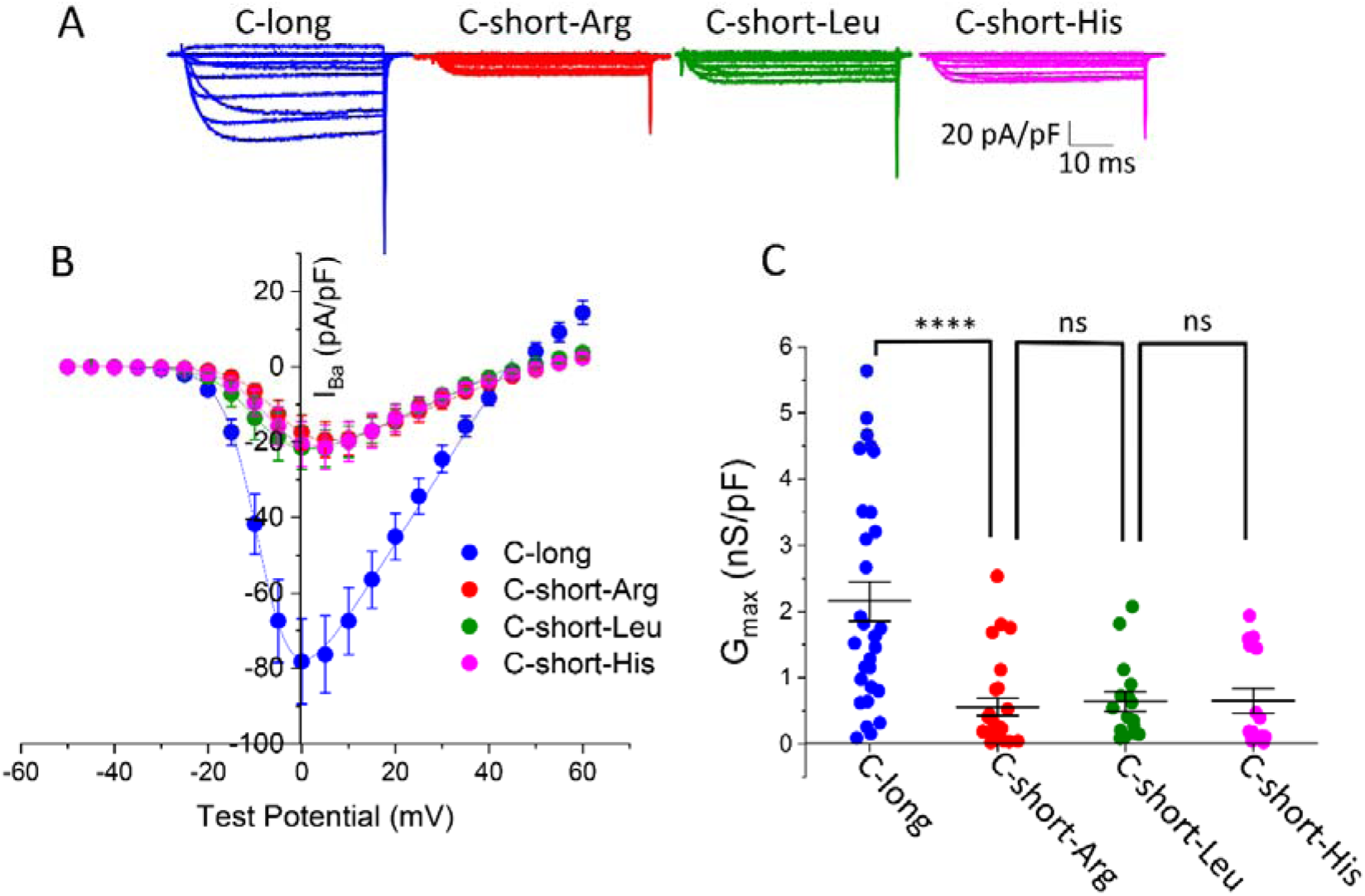
Non-synonymous SNPs in short exon47 do not influence function of CaV2.2. **A** Examples of whole-cell patch-clamp recordings of human GFP_Ca_V_2.2-HA C-long (blue), C-short-Arg (red), C-short-Leu (green) and C-short-His (pink) variants. All conditions are in the presence of α_2_δ-1 and β1b. Holding potential −80 mV, steps between −50 and +60 mV for 50 ms (applies to all traces). **B** Mean (± SEM) *IV* relationships for GFP_Ca_V_2.2-HA, C-long (*n*⍰=⍰30, blue), C-short-Arg (*n*⍰=⍰26, red), C-short-Leu (*n*⍰=⍰16, green) and C-short-His (*n*⍰=⍰18, pink). The individual and mean data were fit with a modified Boltzmann equation (see Materials and Methods). The potential for half-maximal activation (V_50,act_) (mV) was -6.07 ± 0.76, -4.1 ± 0.73, -5.4 ± 1.21 and -5.56 ± 1.13 for C-long, C-short-Arg, C-short-Leu and C-short-His variants respectively. **C** G_max_ (nS/pF) from the *IV* relationships shown in **B**. GFP_Ca_V_2.2-HA C-long and C-short-Arg conditions data are replotted from Fig. 2 for comparison. Individual data (same symbols as in **B**) and mean⍰±⍰SEM are plotted. Statistical significance was determined using one-way ANOVA followed by Šidák *post hoc* test for multiple comparisons: \*\*\*\**P*⍰<⍰0.0001, ns: non-significant.

## Discussion

N-type calcium channels (Ca_V_2.2) are important in primary afferent transmission, including at nociceptor terminals in the dorsal horn. For this reason they are a drug target in pain therapy ^13,14,25-27^. It is therefore particularly important to understand their trafficking and function. All Ca_V_ channels and their auxiliary subunits, like most proteins, have multiple isoforms, conferred by one or more alternatively-spliced or cassette exons, each of which may be differentially expressed in diverse tissues and developmental stages ^18,28,29^. It is therefore essential to understand which combinations of exons are actually included in transcripts in different conditions and tissues ^30,31^, and to examine their combined functional effects. For example, for *CACNA1C* encoding Ca_V_1.2, a large number of novel full length transcript isoforms, many of which are predicted to be functional, were recently identified by long range sequencing in human brain ^32^.

With respect to the multiple alternatively-spliced exons in Ca_V_2.2, several studies have previously examined the effects of individual alternative splicing events ^29^, such as inclusion or skipping of exon18a ^18^, and mutually exclusive exons 37a and 37b ^23,25,33^. In the present study, our initial rationale for examining the effect of the C-terminal short exon47 related to a GWAS study which reported that non-synonymous SNPs in alternatively-spliced exon47 were associated with schizophrenia and Parkinson’s disease ^16^. Because we noted that all reported full length human Ca_V_2.2 transcripts which include short exon47 were missing exon18a, we first examined the effect of inclusion of the long or short exon47 variants on human Ca_V_2.2 function and cell surface expression, in the presence or absence of exon18a. Exon18a encodes 21 residues which are inserted at the proximal end of the intracellular II-III linker. Distal to this region, the II-III linker contains a domain that binds certain synaptic proteins including syntaxin and synaptotagmin ^34,35^, although the importance of this “synprint” domain for presynaptic targeting of calcium channels and for neurotransmitter release is still unclear ^36,37^.

We found that Ca_V_2.2 containing the long exon47, in conjunction with exon18a, supports a large increase in Ca_V_2.2 current amplitude (∼ 4-fold), compared to when exon18a is skipped, or compared to Ca_V_2.2 containing short exon47. In contrast, for Ca_V_2.2 containing short exon47, the inclusion of exon 18a has no effect on calcium current amplitude. Surprisingly, this result is not mirrored by a similar pattern of changes in cell surface expression. Here, the presence of long exon47 supports an ∼4-fold increase in Ca_V_2.2 cell surface trafficking in both tsA-201 cells and in hippocampal neurons, compared to Ca_V_2.2 containing short exon47, irrespective of the presence or absence of exon18. In parallel there is an ∼2-fold increase in the full-length Ca_V_2.2 protein level, conferred by long exon47, as determined by immunoblotting (Figure 3 and Supplementary Figure 2). This result suggests the possibility that if the channel does not reach the cell surface, or is not anchored there, it may be diverted to a degradation pathway, which would be the case for Ca_V_2.2 constructs containing short exon47. In support of this, a lower molecular weight band, which may be a degradation product, is more evident for Ca_V_2.2 containing short exon47 (see Supplementary Figure 2).

Taking these experiments together, it appears that the effect of exon18a inclusion to increase current amplitude, which has been noted previously ^18^, is not mediated primarily by an increase in cell surface expression of the channel. It must rather be mediated by a permissive effect on gating that requires the additional presence of long exon47, since Ca_V_2.2 containing long exon47 mediates a similar increase in cell surface expression irrespective of the presence or absence of exon18a. A corollary of this finding is that the increase in cell surface expression seen for the Δexon18a / long exon47 combination does not result in any increase in Ca_V_2.2 currents, suggesting that the absence of exon18a may exert an inhibitory role for Ca_V_2.2 function. Of great interest, exon18a has been found to be differentially expressed in certain cholecystokinin-containing interneurons ^28^, which depend on N-type Ca channels to mediate GABA release ^38^.

Several previous studies have highlighted the importance of domains in the C-terminus of Ca_V_2.2 in its trafficking and function. In particular, the proximal C-terminus is encoded by exon37, which has two differentially expressed alternative forms, 37a and 37b, that affect Ca_V_2.2 current properties ^33,39^. When exon37a is present, Ca_V_2.2 currents, cell membrane expression and forward trafficking are all increased ^23,33^, via interactions with adaptor protein complex-1 (AP-1) ^23^. All constructs in our study contained exon 37a. Of further relevance to the present study, it has also been found previously that protein-protein interaction motifs in the long C-terminal human Ca_V_2.2 splice variant contain sequences, including a C-terminal Post synaptic density protein, Drosophila disc large tumor suppressor and Zonula occludens-1 protein (PDZ) motif, that are important for its synaptic targeting in cultured hippocampal neurons ^40^. In an attempt to mimic the short human C-terminal Ca_V_2.2, a hybrid human-rat short C-terminal construct was also generated in that study ^40^, although the rat sequence is widely divergent from the human short C-terminus (see Supplementary Figure 3 for comparison), and no full-length rat transcripts containing this short C-terminal sequence have been reported to date. This hybrid construct was found to be mainly concentrated in hippocampal somata and did not extend into neuronal processes ^40^. The human Ca 2.2 containing short exon47, unlike long exon47, does not contain a PDZ motif or SRC Homology 3 (SH3) binding domain (see Supplementary Figure 1). The proline-rich SH3-binding domain sequence, present in long exon47, was identified previously to bind one of the RIM binding protein SH3 domains ^41^. However, in our study a reduction in cell surface expression and trafficking was observed in the non-neuronal tsA-201 cells to the same extent as in hippocampal neuron cell bodies and neurites, indicating that the effect of protein-protein interaction domains in long exon47 is a more general one to enhance cell surface trafficking or anchoring in the cell membrane, rather than (or in addition to) any specific effect to enhance synaptic targeting in neurons.

In this study we observed no effect on Ca_V_2.2 function, expression and trafficking of the SNP variants in short exon47, which encode three different amino acids. This indicates that the observed association of these variants with Parkinson’s disease and schizophrenia ^16^ is not mediated by any grossly altered function. Instead, it is possible that the SNP variants might influence the splicing of exon47, and therefore the relative inclusion of the short and long exon47 variants in Ca_V_2.2 transcripts. In this regard, it is of great interest that Ca_V_2.2 channels play an important role in mediating dopamine release in striatum ^42^.

Future studies will be needed to determine the relative expression of the long and short exon47-containing isoforms in different human tissues and disease states, and whether the SNP alleles influence these properties.

## Acknowledgements

We thank Kanchan Chaggar for technical support, and undergraduate students Catherine Hsu, Helen Cochrane and Syamim Norzihan who were involved in preliminary studies. This work was supported by a Wellcome Trust Investigator award to ACD (098360/Z/12/Z).

## Supplementary Figures

**Supplementary Figure 1.**
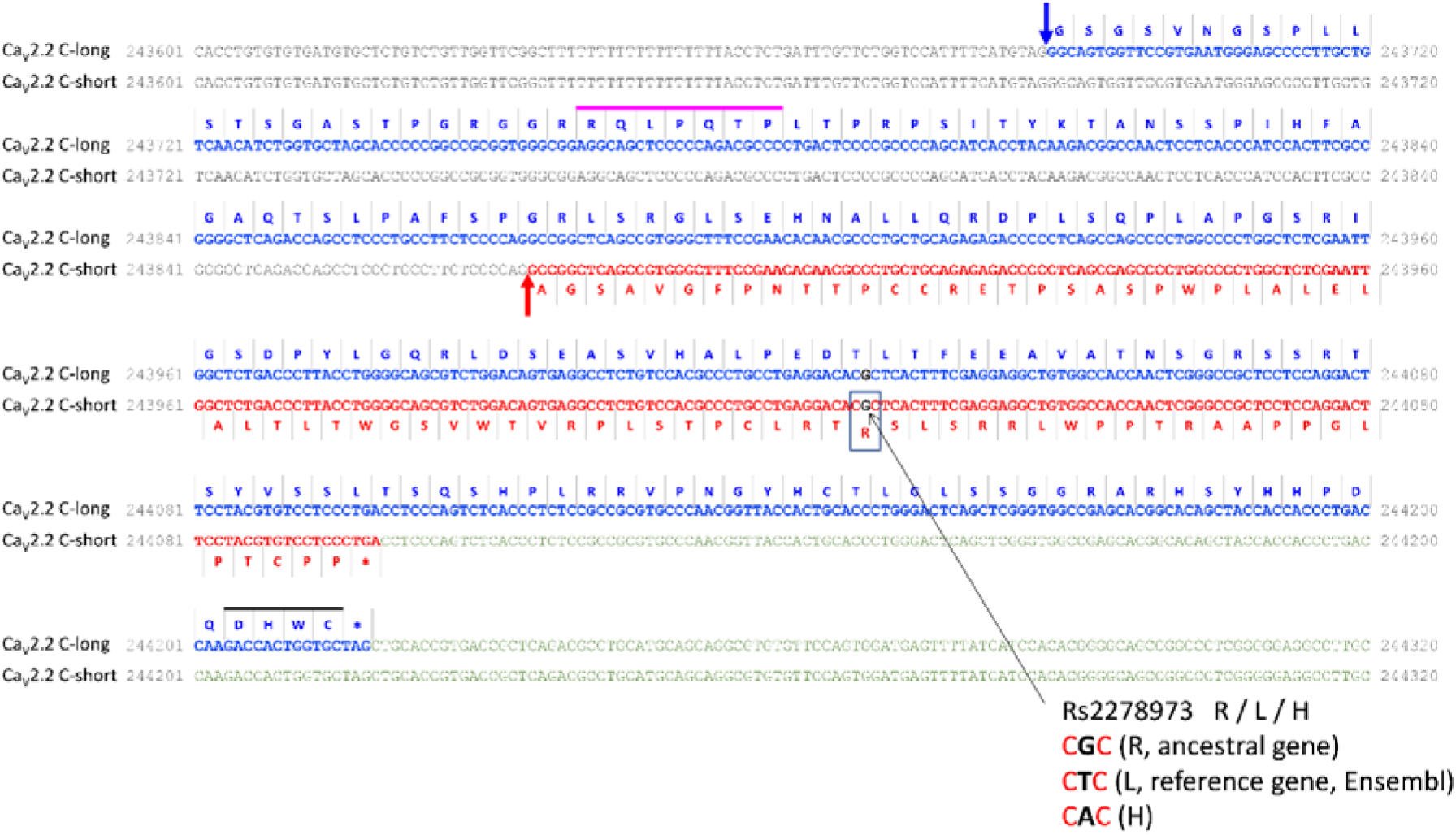
Non-synonymous SNPs in short exon47 of human *CACNA1B*. DNA sequences for the Ca_V_2.2 variants having the long and short C termini. Alternative splice sites are indicated by arrows, blue for the long variant, red for the short. Amino acid sequences are shown in blue for the long variant (above sequence) and red for the short variant (below sequence). Intron sequence is in grey and 3’ untranslated region is in green. The in-frame stop codons for the two variants are indicated by *. The Rs2278973 snp is synonymous in the long variant (ACG/ACT/ACA = T) but results in an amino acid change (CGC=R, CTC=L, CAC=H) in the short variant. According to the gnomAD genomes v3.1.2 database (*Ensembl*), the variants have the following world-wide frequency count: Arg (CGC): 91.7%, Leu (CTC): 8.27%, His (CAC): 0.003%. The proline-rich SH3-binding domain (which bind SH3 domains) was identified previously to bind RIM binding protein ^41^ and is shown as a pink line above the sequence; the PDZ-interacting domain, DXWC ^40,43^ at the end of the C-long sequence, is shown as a black line.

**Supplementary Figure 2.**
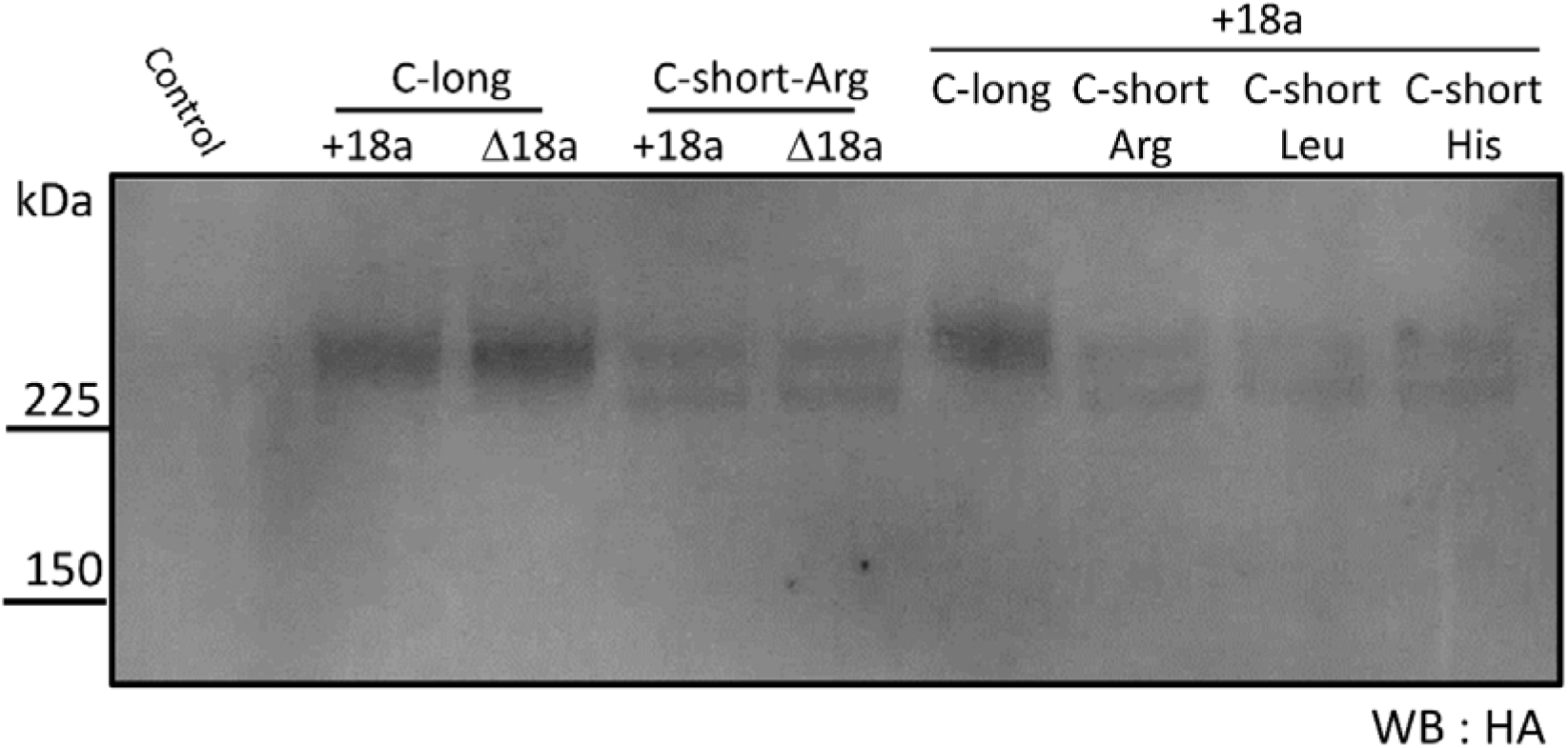
Lack of effect of Ca_V_2.2 C-short SNP variants on total Ca_V_2.2 protein expression. Full immunoblot of whole-cell lysates (WCL) from tsA-201 cells transfected, as stated, with either GFP_Ca_V_2.2-HA C-long, C-short (Arg) (with or without exon18a), C-short-Leu or C-short-His (both + exon18a), together with α_2_δ-1 and β1b. Immunoblot was performed using anti-HA antibody. Partial blot shown in Fig. 5D.

**Supplementary Figure 3.**
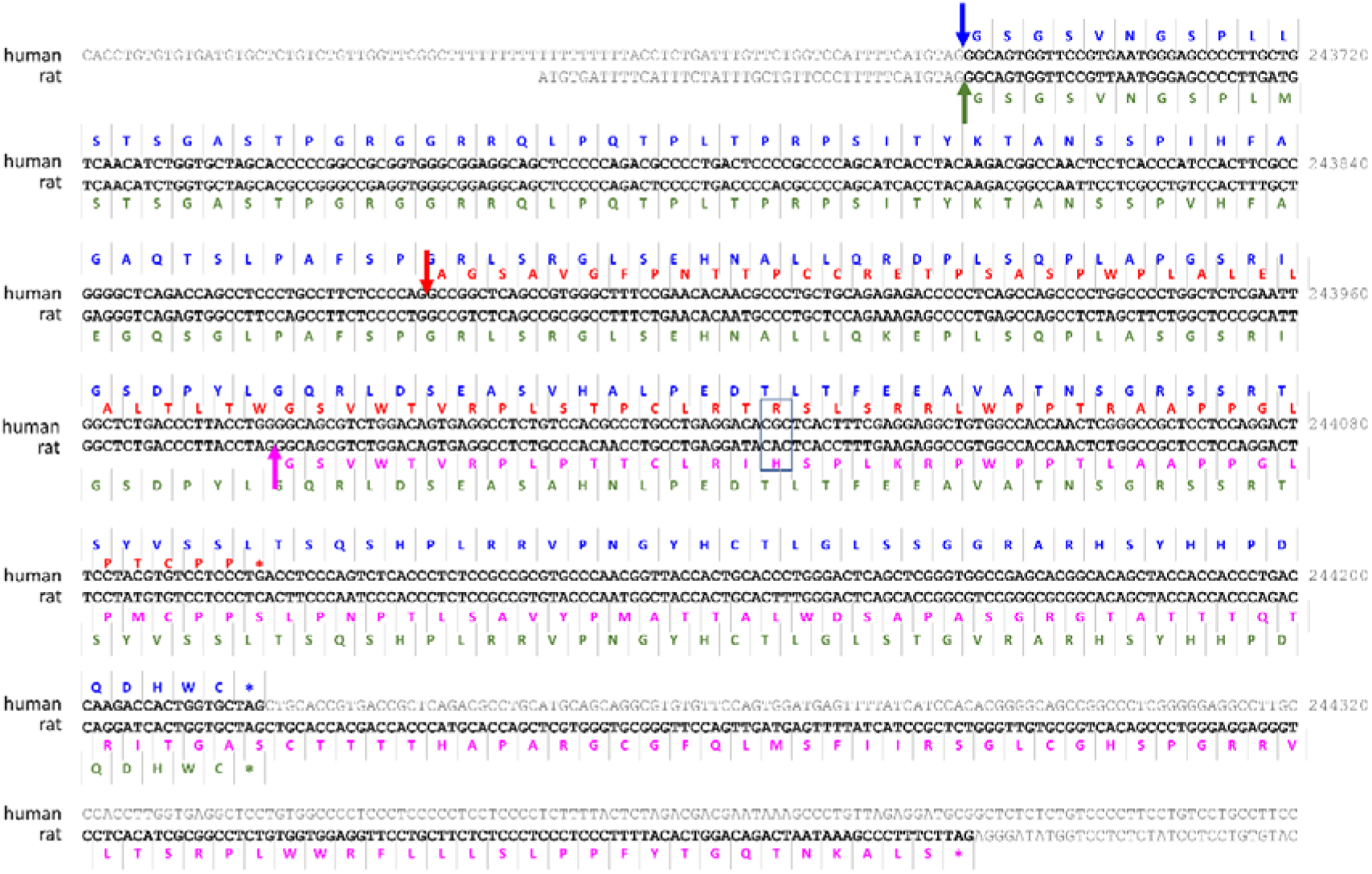
Comparison of organization of rat and human *CACNA1B* transcripts and amino acid sequences. DNA sequences are shown for the Ca_V_2.2 variants having the long and short C termini in human and rat. Alternative splice sites are indicated by arrows, blue for the human long variant, red for the human short, green for the rat long and pink for the rat short. Amino acid sequences are shown in blue for the human long and red for the human short variants (above sequence) and green for the rat long and pink for the rat short variants (below sequence). Intron sequence and 3’ untranslated region is in grey. The in-frame stop codons are indicated by *.

